# Field assessment of precocious maturation in salmon parr using ultrasound imaging

**DOI:** 10.1101/425561

**Authors:** Marie Nevoux, Frédéric Marchand, Guillaume Forget, Dominique Huteau, Julien Tremblay, Jean-Pierre Destouches

**Author notes:** **Cite as:** Nevoux M, Marchand F, Forget G, Huteau D, Tremblay J, and Destouches J-P. Field assessment of precocious maturation in salmon parr using ultrasound imaging. bioRxiv 425561, ver. 3 peer-reviewed and recommended by PCI Ecology (2019). DOI: 10.1101/425561.

## Abstract

Salmonids are characterized by a large diversity of life histories, but their study is often limited by the imperfect observation of the true state of an individual in the wild. Challenged by the need to reduce uncertainty of empirical data, recent development in medical imaging techniques offered new opportunities to assess precocious maturation in Atlantic salmon parr. Traditional phenotypic (external) examination and ultrasound (internal) examination were compared and recommendations on fish handling and ultrasound image interpretation are provided. By allowing to see the unseen, portable ultrasound imaging offers great opportunities for ecological studies in the wild, such as the assessment of individual sexual maturation.

## Introduction

Long term monitoring of animal populations in the wild is of great interest for ecology and management (Clutton-Brock and Sheldon 2010). In a context of global change, such long term monitoring programs contribute to highlight major changes in population composition, dynamics and abundance over the last decades (Parmesan 2006). In the North Pacific, analyses confirmed climate-related shifts in the abundance of most salmonid species since the 1940’s, associated with reported ecosystem regime shifts (Irvine and Fukuwaka 2011). In the North Atlantic, the widespread decline in salmon abundance was associated with a decrease in both survival and age at maturity (Chaput 2012). This global change makes it particularly important to be able to predict population structure and abundance in the future under different climatic scenarios. However, predictive population dynamics models are difficult to parameterize and often return high level of uncertainty, which hinder their use by managers and policy makers. Stock assessments form the basis for catch advice for salmon fisheries, but they mask regional differences and annual river-specific stock assessments are only available for some 25% of the rivers (Chaput 2012). This is partly because field records are imperfect observation of the true state of an individual (Genovart et al. 2012). For instance, maturation status and gender is rarely known with certainty in fish monitoring programs without invasive or lethal methods. Thus, field studies are challenged by the need to reduce uncertainty in individual-level data.

Atlantic salmon (*Salmo salar*) is an anadromous species that reproduces in freshwater and migrates at sea to grow and undergo sexual maturation. However, in this and other salmonid species, some young males (parrs) can mature precociously in freshwater (Fleming 1996). It is proposed that the intensity of intra-sexual competition for mates and environmental variability in growth opportunities, determines a polygenic threshold that triggers precocious maturation (Hutchings and Myers 1994). Mature male parrs attempt to ‘sneak’ access to large anadromous females for mating (Fleming 1996). They can contribute substantially to the reproduction (Moran et al. 1996) but, as maturation seems to be traded against survival, mature parr have a low probability to migrate at sea (Buoro et al. 2010). The occurrence of this alternative reproduction strategy is highly variable over time, space, and age classes (Baglinière and Maisse 1985; Myers et al. 1986) and may be driven by frequency dependent selection, i.e. the reproductive success of precocious parrs increases with decreasing frequency of precocious maturation in the population (Berejikian et al. 2010). Yet, precise quantification of this phenotype remains rare in empirical studies and in salmon population dynamic models. Furthermore, precocious maturation is virtually ignored by stock assessment models and in the management of Atlantic salmon fisheries (e.g. ICES 2013). This means that mature parrs are included neither in the stock nor in the recruitment figures. This might bias stock-recruitment relationships and underestimate freshwater survival as well as salmon production. For instance, using more than 280,000 records from Little Codroy River parrs, Myers (1984) estimated that 60% of the stock of adult male salmon in Newfoundland was lost because of increased mortality due to precocious maturation. Caswell et al. (1984) demonstrated that the effect of reproduction by parrs on population growth rate was always greater than that of reproduction by adults. It is suspected that this lack of consideration for mature parrs is due to the complexity of the life cycle of Atlantic salmon. But it could also be due to the difficulty of quantifying the occurrence of precocious maturation in wild salmon populations.

In the field, precocious maturity in parrs is traditionally assessed by pressing the abdomen of the fish to extract milt (sperm). However, not all mature males may be producing milt at the time of capture. Moreover, based on the general aspect of the fish, experienced fieldworkers sometimes suspect maturity in some individuals that do not excrete milt upon abdominal pressure. However, researchers are often reluctant to use such expert opinions in their analysis due to the potential for incorrect diagnosis and strong operator bias.

Ultrasound imaging, which gave rise to significant advances in human medicine, offers reliable visualization of internal organs. Ultrasound imaging is routinely used in aquaculture as it provides an accurate, non-invasive, non-lethal method that is more accurate that visual methods, to determine gender or gonadal growth in fish for example (Novelo and Tiersch 2012). In adult salmon, although gonads of immature males may be difficult to discern, gonads of females are always visible and hence sexing is possible by deduction (Martin et al. 1983; Mattson 1991). However the ovaries and testes of the juvenile (immature) salmons seem difficult to identify (Martin et al. 1983).

The development of portable devices offers an opportunity to bring ultrasound machines in the field. This paper illustrates how ultrasound scanners can increase the range of biological traits that can be monitored in long-term salmonid populations surveys. In particular, the maturation status of salmon parr (i.e. freshwater resident salmon) was assessed in the field using traditional phenotypic examination and ultrasound examination. The two approaches are compared and it is tested whether biological or operator-related factors affect the uncertainty in external phenotypic assessment of parr maturity. Finally, recommendation on fish handling and on the interpretation of ultrasound images is provided.

## Methods

### Salmon monitoring

Atlantic salmon monitoring takes place every year in September or October in the river Oir, Normandy, France (Marchand et al. 2018). Young individuals (parrs) are captured using a standard electrofishing protocol. They are then placed in a light anesthetic solution, measured, weighed, scanned, and returned to the river within 30 min of their capture. Age is assessed through scale reading. In 2015, 2016, and 2017, a total of 850 salmon was examined (366 parrs of age 0 and 484 parrs of age 1) for maturity using a traditional, phenotypic (i.e. external), approach and ultrasound imaging. All parrs were caught within a period of 8 days in early October. All the animal experimentation in this study was performed according to French legislation and under licence APAFIS-201602051204637. The data is available in Appendix 1.

### Traditional phenotypic examination

First, the field operator assesses the maturation state of parrs by gently pressing the side of the fish. The excretion of milt is indicative of a mature male, while the absence of milt can be encountered in a maturing male whose gonads do not yet produce milt, an immature male, or a female (females are all immature at this stage). The field operator may also report that some fish *look like* mature parrs: they display phenotypic traits similar to those of mature fish (e.g. colour pattern, body shape) but do not excrete milt. This phenotypic assessment relies less on a fixed set of quantifiable parameters than on a general, and subjective, appreciation of the fish, which is specific to each operator.

### Ultrasound examination

Then, the ultrasound operator screens all salmon parrs for precocious maturation using a portable ultrasound scanner M-Turbo (Sonosite) with a 5-10 MHz linear transducer. The default setting, for muscular examination, is selected. As water perfectly transmits ultrasounds, the use of ultrasound transmission gel is not needed. Fish are placed on their side in a tank of freshwater and the transducer is operated in the water, keeping it 1-2cm above the belly of the fish. Sagittal images of the abdominal cavity are produced, i.e. the transducer is aligned with the lateral line. The maturation state of each individual is directly assessed from live images on the ultrasound monitor. Parrs are classified as mature when gonads are detected, or immature when the gonads are too small to be detected. Ultrasound snapshots can be stored on the ultrasound machine and then exported to a USB memory stick for post-processing.

### Comparison of phenotypic and ultrasound methods

The comparison of the proportion of parrs that are recorded each year as mature or immature by each approach is done with a Chi² test. The number of mature parrs is defined as the sum of individuals whose gonads are detected with the ultrasound and individuals whose gonads are not detected with the ultrasound but that produce milt. Because the probability to detect precocious maturation with the phenotypic method depends on the probability for a mature male to produce milt at the time of capture, an analysis is conducted to investigate which biological factor drive this probability. Empirical records show a large interannual variability in the proportion of maturing parrs (Baglinière and Maisse 1985; Myers et al. 1986). Similarly, the year of capture is expected to affect milting within mature parrs. Thus, a generalized linear model (GLM) with a binomial distribution of errors is used to test for a year effect. Then, a mixed model with year as a random intercept and a binomial distribution of errors is used to test for the effect of age, body length, and their interaction. A GLM is also run to test whether the probability to miss a mature parr that does not produce milt is affected by age, body length, operator identity, and time of day (testing for operator tiredness). The date of capture is not considered because of the low variance in this variable. The significance of the above-mentioned effects is assessed using the z-value. All models are run in R (R Development Core Team 2018).

## Results and discussion

### Ultrasound assessment of precocious maturity

A trained ultrasound operator conducts ultrasound examination for maturation assessment in ca. 5 seconds. In immature individuals, the liver, stomach, caeca, and intestine can be identified (Figure 1a). Note that all internal organs appear more clearly on live images, during ultrasound examination, than on the snapshots displayed here. Nevertheless, the heterogeneous filling and uneven granularity of the abdominal cavity are diagnostic of immature individuals, as well as the concave shape of the cavity, which appears depressed on both sides. In mature parrs (precocious males), gonads virtually fill the whole abdominal cavity and ultrasound snapshots depict a homogeneous and finely granulated pattern (Figure 1b, c); digestive organs are hardly visible and the cavity is convex. In mature males that do not produce milt (Figure 1b), gonads appear less developed and fill the cavity to a lesser extent than in males producing milt (Figure 1c), which is consistent with a lower degree of maturity in the former. Interestingly, ultrasound imaging provides the same objective diagnosis of maturation in males that do not produce milt (Figure 1b) and ones that do (Figure 1c). It also discriminates truly mature parrs from well-fed individuals, which the operator can mistake for mature males upon visual inspection because of their round belly.

**Figure 1.**
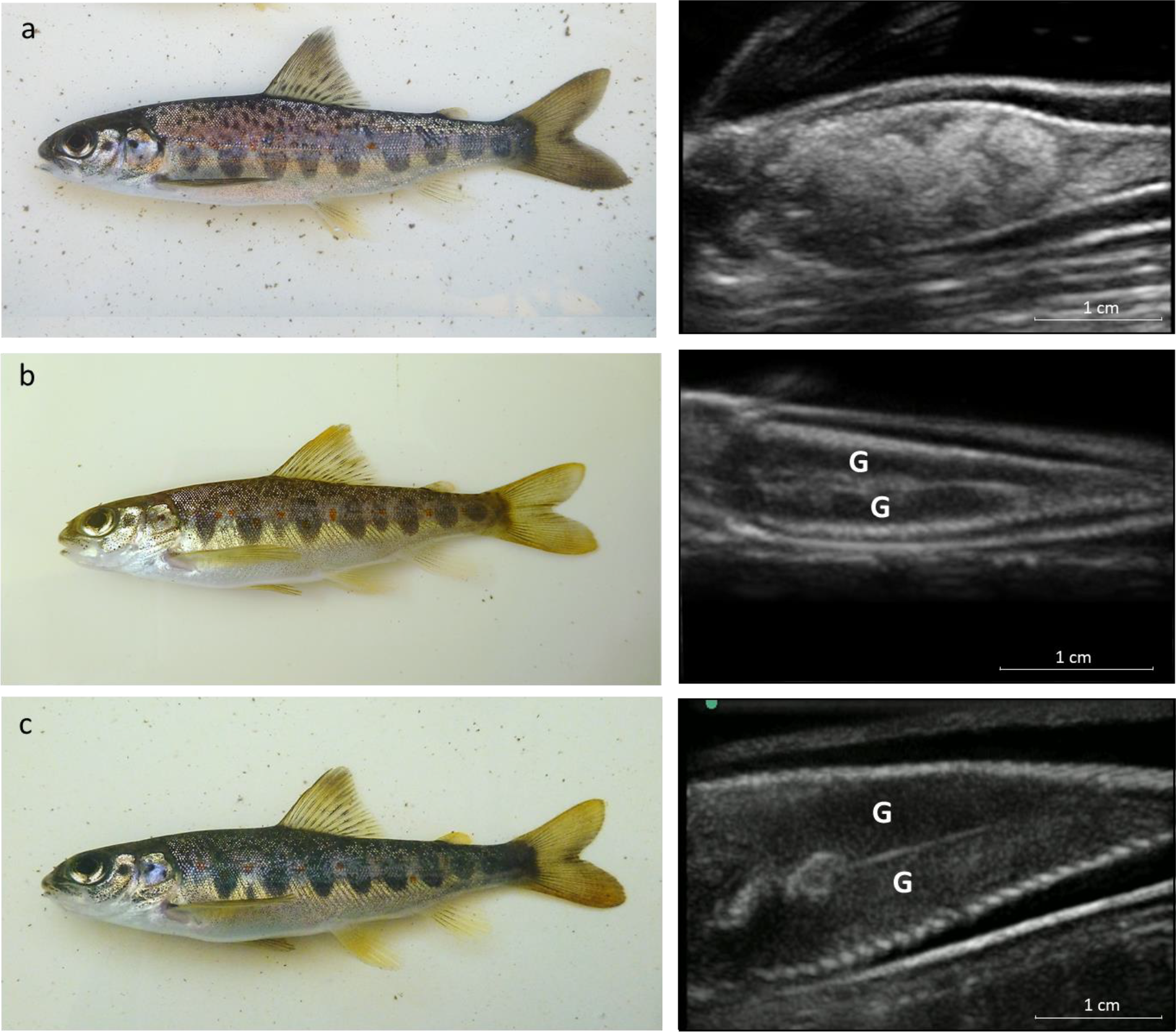
Assessing maturity in salmon parr using phenotypic view (left panel) and ultrasound imaging (right panel): a) immature one-year-old parr, b) mature young-of-the-year parr that does not produce milt (precocious male), and c) mature one-year-old parr that produces milt (precocious male) in river Oir. On ultrasound snapshots, the head of the fish is on the left but not shown, the left flank is towards the bottom of the snapshot and the right towards the top. Gonads are identified by the letter “G”.

### Comparison of phenotypic and ultrasound methods

The proportion of mature parrs detected by the phenotypic method is much lower and more variable (63.8 %, SD = 40.7%) than with the ultrasound method (95.4%, SD = 7.1%, p = 0.199). On average 33.4% of mature males do not produce milt. A strong year effect is detected (p < 0.001) in the probability to produce milt at the time of capture in mature males. When accounting for this year effect as a random factor, a strong increase with age (estimate = 5.582, SE = 1.031, p < 0.001) and a negative interaction with body length within each age class (estimate = −0.031, SE = 0.008, p < 0.001) are detected. This means that a higher proportion of mature males produce milt in autumn in old parrs than in young ones and, within a given age class in small parrs than in large ones.

Because the probability to detect precocious maturation with the phenotypic method depends on the probability for a mature male to produce milt at the time of capture, our result highlights that both intrinsic and environmental factors can affect the level of detection of the precocious state. The ultrasound method shows that only 16.8% and 13.2% of the mature males are identified as precocious parrs at the time of capture in 2015 and 2017, while this proportion rises to 87.6% in 2016 (Table 1). Empirical evidence have shown an increase in the proportion of mature parrs as the parrs are getting older (Baglinière and Maisse 1985; Myers et al. 1986). But this study suggests that mature parrs develop and produce milt earlier in the season than young ones, which may accentuate even further the age-specific pattern detected above, with the phenotypic method.

**Table 1.** Comparison of the number of records using external phenotypic observation (rows) and ultrasound imaging (columns) to assess maturation in parrs each year.

| Year | Phenotype method \ ultrasound method | Gonads detected with ultrasound | Gonads not detected with ultrasound |
| --- | --- | --- | --- |
| 2015 | Parr producing milt | 17 | 0 |
|  | Parr not producing milt that looks like a mature parr | 74 | 8 |
|  | Parr not producing milt that looks like an immature parr | 10 | 73 |
| 2016 | Parr producing milt | 181 | 2 |
|  | Parr not producing milt that looks like a mature parr | 20 | 3 |
|  | Parr not producing milt that looks like an immature parr | 6 | 251 |
| 2017 | Parr producing milt | 23 | 4 |
|  | Parr not producing milt that looks like a mature parr | 1 | 0 |
|  | Parr not producing milt that looks like an immature parr | 3 | 174 |

In our study, the maturation state of 106 individuals is uncertain according to the phenotypic method, i.e. individuals that look like mature parrs but do not produce milt. However, following ultrasound examination, 89.6 % of those parrs are indeed found to be mature. This result acknowledges the high level of expertise of our trained operators to assess the maturation state of parrs from external phenotypic examination. Conversely, gonads are not detected in 10.4% of those parrs (false positive) and the field operators do not detect milt nor suspect maturity in 5.6 % of the actually mature males (false negative). This later proportion differs between the two operators (p = 0.008). None of the other variables has a significant effect. This highlights how difficult it is to assess parr maturation from external examination in the field in a fully reliable and replicable manner.

Contrary to the phenotypic method, it is assumed that the ultrasound method cannot produce false positives, because gonads large enough to be detected by the ultrasound equipment are only found in mature parrs. The ultrasound operator missed 2.7% of the males producing milt (false negative). The sample size is too small to assess the potential effect of external factors on this proportion through multivariate analysis. Still, this shows that ultrasound imaging is still subject to a low degree of uncertainty in maturity assessment. The correct interpretation of ultrasound images depends on the skill of the operator. Indeed, some training is required to operate the transducer in a fluent way and navigate between different anatomic layers. In this regard, a good knowledge of the internal structure for the species of interest is a prerequisite. This improves the quality of the ultrasound snapshots and minimizes misinterpretation issues. In hindsight, it is also recommended that all ultrasounds should be examined and interpreted by multiple operators. This ensures a quick examination and thus minimizes the impact on fish wellbeing.

### Conclusion

Assessing precocious maturation in parrs from phenotypic observation can reach a high level of accuracy in trained operators. However, this method relies on some subjective criteria and gut feeling, which render the level of accuracy in the data difficult to quantify. By allowing to see the unseen, portable ultrasound imaging offers great opportunities for ecological studies in the wild. As illustrated in this paper, key phenotypic traits become accessible and help better characterize the true biological state of individuals. Ultrasound imaging is an objective, easily transferable and replicable approach to investigate precocious maturity in parrs, as it relies on key diagnostic features. A naïve operator can gain a good expertise and assess precocious maturation with the ultrasound method within a day, whereas more training would be required to achieve a similar level of expertise with the phenotypic method. In the end, this study calls for the assessment of precocious maturation in Atlantic salmon parrs using the two methods together: 1) testing whether parrs produce milt is a quick examination and provides an objective and undeniable evidence of maturity, 2) ultrasound imaging should remove any ambiguity about the state of maturation in parrs that do not produce milt. Reducing uncertainty in empirical data this way should offer new opportunities for further research on this alternative breeding strategy in salmon and improve our understanding of this key biological process.

## Acknowledgements

This preprint has been peer-reviewed and recommended by Peer Community In Ecology (https://dx.doi.org/10.24072/pci.ecology.100021).

## Appendix

## Appendix 1

Individual records of date and hour of capture, body size, age, phenotypic observation and ultrasound examination for the 850 salmon parrs captured in the river Oir for this study.

| Date | Hour | Size (mm) | Age | Phenotype maturation | Phenotype operator | Ultrasound maturation |
| --- | --- | --- | --- | --- | --- | --- |
| 05/10/2015 | 10 | 95 | 1 | milt | FM | mature |
| 05/10/2015 | 10 | 115 | 1 | milt | FM | mature |
| 05/10/2015 | 10 | 102 | 1 | milt | FM | mature |
| 05/10/2015 | 10 | 113 | 1 | milt | FM | mature |
| 05/10/2015 | 10 | 125 | 1 | milt | FM | mature |
| 05/10/2015 | 10 | 90 | 1 | milt | FM | mature |
| 05/10/2015 | 10 | 111 | 1 | milt | FM | mature |
| 05/10/2015 | 10 | 104 | 1 | milt | FM | mature |
| 05/10/2015 | 10 | 105 | 1 | milt | FM | mature |
| 05/10/2015 | 10 | 117 | 1 | milt | FM | mature |
| 05/10/2015 | 10 | 105 | 1 | milt | FM | mature |
| 05/10/2015 | 11 | 120 | 1 | milt | FM | mature |
| 05/10/2015 | 11 | 109 | 1 | milt | FM | mature |
| 05/10/2015 | 11 | 111 | 1 | immature | FM | immature |
| 05/10/2015 | 11 | 110 | 1 | immature | FM | immature |
| 05/10/2015 | 11 | 116 | 1 | immature | FM | immature |
| 05/10/2015 | 11 | 124 | 1 | immature | FM | mature |
| 05/10/2015 | 11 | 117 | 1 | immature | FM | immature |
| 05/10/2015 | 11 | 112 | 1 | immature | FM | immature |
| 05/10/2015 | 12 | 79 | 0 | immature | FM | mature |
| 05/10/2015 | 12 | 71 | 0 | mature | FM | mature |
| 05/10/2015 | 12 | 123 | 1 | immature | FM | immature |
| 05/10/2015 | 12 | 121 | 1 | immature | FM | immature |
| 05/10/2015 | 12 | 107 | 1 | immature | FM | immature |
| 05/10/2015 | 12 | 120 | 1 | mature | FM | mature |
| 05/10/2015 | 12 | 130 | 1 | immature | FM | immature |
| 05/10/2015 | 12 | 77 | 0 | mature | FM | mature |
| 05/10/2015 | 12 | 87 | 0 | immature | FM | immature |
| 05/10/2015 | 12 | 75 | 0 | mature | FM | mature |
| 05/10/2015 | 14 | 112 | 1 | immature | FM | immature |
| 05/10/2015 | 13 | 102 | 1 | immature | FM | immature |
| 05/10/2015 | 13 | 112 | 1 | immature | FM | immature |
| 05/10/2015 | 13 | 107 | 1 | mature | FM | mature |
| 05/10/2015 | 14 | 61 | 0 | mature | FM | mature |
| 05/10/2015 | 13 | 100 | 1 | immature | FM | mature |
| 05/10/2015 | 13 | 102 | 1 | immature | FM | immature |
| 05/10/2015 | 14 | 115 | 1 | mature | FM | mature |
| 05/10/2015 | 13 | 93 | 1 | mature | FM | mature |
| 05/10/2015 | 13 | 104 | 1 | immature | FM | mature |
| 05/10/2015 | 13 | 125 | 1 | immature | FM | immature |
| 05/10/2015 | 14 | 110 | 1 | immature | FM | immature |
| 05/10/2015 | 13 | 93 | 1 | mature | FM | mature |
| 05/10/2015 | 13 | 121 | 1 | immature | FM | immature |
| 05/10/2015 | 13 | 108 | 1 | mature | FM | mature |
| 05/10/2015 | 13 | 92 | 1 | mature | FM | mature |
| 05/10/2015 | 13 | 112 | 1 | immature | FM | immature |
| 05/10/2015 | 13 | 100 | 1 | mature | FM | mature |
| 05/10/2015 | 13 | 75 | 0 | mature | FM | mature |
| 05/10/2015 | 15 | 105 | 1 | mature | FM | mature |
| 05/10/2015 | 15 | 121 | 1 | mature | FM | mature |
| 05/10/2015 | 15 | 112 | 1 | immature | FM | immature |
| 05/10/2015 | 15 | 76 | 0 | mature | FM | mature |
| 05/10/2015 | 15 | 100 | 1 | mature | FM | mature |
| 05/10/2015 | 15 | 109 | 1 | mature | FM | mature |
| 05/10/2015 | 15 | 72 | 0 | mature | FM | mature |
| 05/10/2015 | 15 | 108 | 1 | immature | FM | immature |
| 05/10/2015 | 15 | 94 | 1 | mature | FM | mature |
| 05/10/2015 | 15 | 68 | 0 | immature | FM | immature |
| 05/10/2015 | 15 | 103 | 1 | mature | FM | mature |
| 05/10/2015 | 15 | 119 | 1 | mature | FM | mature |
| 05/10/2015 | 15 | 116 | 1 | mature | FM | mature |
| 05/10/2015 | 15 | 71 | 0 | mature | FM | mature |
| 05/10/2015 | 15 | 122 | 1 | immature | FM | immature |
| 05/10/2015 | 15 | 110 | 1 | immature | FM | immature |
| 05/10/2015 | 15 | 110 | 1 | mature | FM | mature |
| 05/10/2015 | 15 | 120 | 1 | immature | FM | immature |
| 05/10/2015 | 16 | 103 | 1 | mature | FM | mature |
| 05/10/2015 | 17 | 115 | 1 | immature | FM | immature |
| 05/10/2015 | 16 | 91 | 1 | milt | FM | mature |
| 05/10/2015 | 16 | 100 | 1 | immature | FM | mature |
| 05/10/2015 | 17 | 66 | 0 | mature | FM | mature |
| 05/10/2015 | 16 | 72 | 0 | immature | FM | mature |
| 05/10/2015 | 16 | 122 | 1 | mature | FM | mature |
| 05/10/2015 | 16 | 127 | 1 | immature | FM | immature |
| 05/10/2015 | 17 | 79 | 0 | mature | FM | immature |
| 05/10/2015 | 16 | 115 | 1 | immature | FM | immature |
| 05/10/2015 | 16 | 101 | 1 | milt | FM | mature |
| 05/10/2015 | 17 | 102 | 1 | immature | FM | immature |
| 05/10/2015 | 17 | 65 | 0 | milt | FM | mature |
| 05/10/2015 | 17 | 122 | 1 | immature | FM | immature |
| 05/10/2015 | 17 | 110 | 1 | immature | FM | immature |
| 05/10/2015 | 17 | 102 | 1 | immature | FM | immature |
| 05/10/2015 | 17 | 103 | 1 | immature | FM | immature |
| 05/10/2015 | 16 | 110 | 1 | immature | FM | immature |
| 05/10/2015 | 16 | 106 | 1 | mature | FM | mature |
| 05/10/2015 | 16 | 71 | 0 | mature | FM | immature |
| 05/10/2015 | 17 | 70 | 0 | immature | FM | immature |
| 06/10/2015 | 10 | 99 | 1 | immature | FM | mature |
| 06/10/2015 | 10 | 117 | 1 | immature | FM | immature |
| 06/10/2015 | 11 | 111 | 1 | immature | FM | immature |
| 06/10/2015 | 11 | 119 | 1 | immature | FM | immature |
| 06/10/2015 | 10 | 102 | 1 | immature | FM | immature |
| 06/10/2015 | 10 | 115 | 1 | mature | FM | mature |
| 06/10/2015 | 10 | 125 | 1 | mature | FM | mature |
| 06/10/2015 | 10 | 106 | 1 | mature | FM | mature |
| 06/10/2015 | 10 | 100 | 1 | mature | FM | mature |
| 06/10/2015 | 09 | 110 | 1 | mature | FM | mature |
| 06/10/2015 | 10 | 70 | 0 | mature | FM | mature |
| 06/10/2015 | 10 | 116 | 1 | immature | FM | immature |
| 06/10/2015 | 10 | 123 | 1 | immature | FM | immature |
| 06/10/2015 | 10 | 71 | 0 | immature | FM | immature |
| 06/10/2015 | 10 | 110 | 1 | mature | FM | mature |
| 06/10/2015 | 10 | 97 | 1 | mature | FM | mature |
| 06/10/2015 | 10 | 109 | 1 | mature | FM | mature |
| 06/10/2015 | 11 | 84 | 0 | immature | FM | mature |
| 06/10/2015 | 12 | 104 | 1 | immature | FM | immature |
| 06/10/2015 | 12 | 79 | 0 | mature | FM | mature |
| 06/10/2015 | 11 | 125 | 1 | immature | FM | immature |
| 06/10/2015 | 11 | 109 | 1 | immature | FM | immature |
| 06/10/2015 | 12 | 105 | 1 | immature | FM | mature |
| 06/10/2015 | 12 | 72 | 0 | mature | FM | mature |
| 06/10/2015 | 12 | 112 | 1 | immature | FM | immature |
| 06/10/2015 | 11 | 113 | 1 | immature | FM | immature |
| 06/10/2015 | 14 | 110 | 1 | immature | FM | immature |
| 06/10/2015 | 15 | 138 | 1 | immature | FM | immature |
| 06/10/2015 | 15 | 119 | 1 | immature | FM | immature |
| 06/10/2015 | 15 | 112 | 1 | mature | FM | mature |
| 06/10/2015 | 14 | 110 | 1 | immature | FM | immature |
| 06/10/2015 | 13 | 118 | 1 | immature | FM | immature |
| 06/10/2015 | 14 | 120 | 1 | immature | FM | immature |
| 06/10/2015 | 13 | 115 | 1 | immature | FM | immature |
| 06/10/2015 | 15 | 115 | 1 | immature | FM | immature |
| 06/10/2015 | 16 | 65 | 0 | mature | FM | mature |
| 06/10/2015 | 15 | 112 | 1 | immature | FM | immature |
| 06/10/2015 | 15 | 110 | 1 | immature | FM | immature |
| 06/10/2015 | 16 | 80 | 0 | immature | FM | immature |
| 06/10/2015 | 15 | 74 | 0 | mature | FM | mature |
| 06/10/2015 | 15 | 69 | 0 | mature | FM | immature |
| 06/10/2015 | 15 | 97 | 1 | immature | FM | immature |
| 06/10/2015 | 15 | 114 | 1 | mature | FM | mature |
| 06/10/2015 | 15 | 127 | 1 | immature | FM | immature |
| 06/10/2015 | 16 | 106 | 1 | mature | FM | mature |
| 06/10/2015 | 17 | 106 | 1 | mature | FM | mature |
| 06/10/2015 | 18 | 102 | 1 | mature | FM | mature |
| 06/10/2015 | 16 | 87 | 0 | immature | FM | immature |
| 06/10/2015 | 17 | 121 | 1 | mature | FM | mature |
| 06/10/2015 | 16 | 105 | 1 | immature | FM | immature |
| 06/10/2015 | 17 | 108 | 1 | immature | FM | immature |
| 06/10/2015 | 17 | 103 | 1 | immature | FM | immature |
| 06/10/2015 | 17 | 122 | 1 | immature | FM | immature |
| 06/10/2015 | 16 | 107 | 1 | immature | FM | immature |
| 06/10/2015 | 17 | 108 | 1 | immature | FM | immature |
| 06/10/2015 | 17 | 105 | 1 | mature | FM | mature |
| 06/10/2015 | 17 | 105 | 1 | immature | FM | immature |
| 06/10/2015 | 17 | 121 | 1 | immature | FM | immature |
| 06/10/2015 | 17 | 113 | 1 | immature | FM | immature |
| 06/10/2015 | 17 | 104 | 1 | immature | FM | immature |
| 06/10/2015 | 17 | 72 | 0 | mature | FM | mature |
| 06/10/2015 | 10 | 94 | 1 | immature | FM | immature |
| 06/10/2015 | 18 | 102 | 1 | immature | FM | immature |
| 06/10/2015 | 10 | 97 | 1 | mature | FM | mature |
| 07/10/2015 | 10 | 122 | 1 | mature | FM | immature |
| 07/10/2015 | 11 | 70 | 0 | mature | FM | mature |
| 07/10/2015 | 11 | 72 | 0 | mature | FM | mature |
| 07/10/2015 | 11 | 108 | 1 | mature | FM | immature |
| 07/10/2015 | 11 | 77 | 0 | mature | FM | mature |
| 07/10/2015 | 12 | 113 | 1 | mature | FM | mature |
| 07/10/2015 | 12 | 121 | 1 | mature | FM | immature |
| 07/10/2015 | 12 | 81 | 0 | mature | FM | mature |
| 07/10/2015 | 14 | 111 | 1 | mature | FM | mature |
| 07/10/2015 | 14 | 101 | 1 | mature | FM | mature |
| 07/10/2015 | 13 | 100 | 1 | mature | FM | mature |
| 07/10/2015 | 14 | 110 | 1 | mature | FM | mature |
| 07/10/2015 | 16 | 71 | 0 | mature | FM | mature |
| 08/10/2015 | 10 | 115 | 1 | mature | FM | mature |
| 08/10/2015 | 09 | 78 | 0 | mature | FM | mature |
| 08/10/2015 | 11 | 75 | 0 | mature | FM | mature |
| 08/10/2015 | 11 | 133 | 1 | mature | FM | mature |
| 08/10/2015 | 09 | 89 | 1 | mature | FM | mature |
| 08/10/2015 | 09 | 100 | 1 | immature | FM | mature |
| 08/10/2015 | 09 | 80 | 0 | mature | FM | mature |
| 08/10/2015 | 10 | 124 | 1 | mature | FM | immature |
| 08/10/2015 | 10 | 118 | 1 | milt | FM | mature |
| 08/10/2015 | 10 | 73 | 0 | mature | FM | mature |
| 08/10/2015 | 11 | 67 | 0 | immature | FM | immature |
| 08/10/2015 | 12 | 81 | 0 | mature | FM | mature |
| 08/10/2015 | 11 | 125 | 1 | mature | FM | mature |
| 08/10/2015 | 12 | 114 | 1 | mature | FM | mature |
| 08/10/2015 | 13 | 122 | 1 | mature | FM | mature |
| 08/10/2015 | 12 | 82 | 0 | mature | FM | immature |
| 08/10/2015 | 12 | 100 | 1 | mature | FM | mature |
| 08/10/2015 | 13 | 72 | 0 | mature | FM | mature |
| 03/10/2016 | 08 | 116 | 1 | milt | FM | mature |
| 03/10/2016 | 08 | 113 | 1 | milt | FM | mature |
| 03/10/2016 | 08 | 106 | 1 | immature | FM | immature |
| 03/10/2016 | 08 | 72 | 0 | immature | FM | immature |
| 03/10/2016 | 08 | 77 | 0 | immature | FM | immature |
| 03/10/2016 | 08 | 74 | 0 | immature | FM | immature |
| 03/10/2016 | 09 | 125 | 1 | immature | FM | immature |
| 03/10/2016 | 09 | 70 | 0 | immature | FM | immature |
| 03/10/2016 | 09 | 119 | 1 | milt | FM | mature |
| 03/10/2016 | 09 | 113 | 1 | immature | FM | immature |
| 03/10/2016 | 09 | 68 | 0 | immature | FM | immature |
| 03/10/2016 | 09 | 100 | 1 | milt | FM | mature |
| 03/10/2016 | 09 | 120 | 1 | immature | FM | immature |
| 03/10/2016 | 09 | 113 | 1 | immature | FM | immature |
| 03/10/2016 | 09 | 69 | 0 | immature | FM | immature |
| 03/10/2016 | 09 | 105 | 1 | immature | FM | immature |
| 03/10/2016 | 09 | 74 | 0 | immature | FM | immature |
| 03/10/2016 | 09 | 128 | 1 | immature | FM | immature |
| 03/10/2016 | 09 | 117 | 1 | milt | FM | mature |
| 03/10/2016 | 09 | 83 | 0 | immature | FM | immature |
| 03/10/2016 | 09 | 108 | 1 | milt | FM | mature |
| 03/10/2016 | 09 | 133 | 1 | milt | FM | mature |
| 03/10/2016 | 09 | 126 | 1 | milt | FM | mature |
| 03/10/2016 | 09 | 118 | 1 | mature | FM | mature |
| 03/10/2016 | 10 | 116 | 1 | milt | FM | mature |
| 03/10/2016 | 10 | 99 | 1 | milt | FM | mature |
| 03/10/2016 | 10 | 72 | 0 | immature | FM | immature |
| 03/10/2016 | 10 | 76 | 0 | milt | FM | mature |
| 03/10/2016 | 10 | 113 | 1 | milt | FM | mature |
| 03/10/2016 | 10 | 97 | 0 | immature | FM | immature |
| 03/10/2016 | 10 | 88 | 0 | immature | FM | immature |
| 03/10/2016 | 10 | 85 | 0 | milt | FM | mature |
| 03/10/2016 | 10 | 69 | 0 | immature | FM | immature |
| 03/10/2016 | 10 | 84 | 0 | immature | FM | immature |
| 03/10/2016 | 10 | 105 | 1 | mature | FM | mature |
| 03/10/2016 | 10 | 115 | 1 | milt | FM | mature |
| 03/10/2016 | 10 | 111 | 1 | milt | FM | mature |
| 03/10/2016 | 10 | 113 | 1 | milt | FM | mature |
| 03/10/2016 | 10 | 119 | 1 | milt | FM | mature |
| 03/10/2016 | 10 | 79 | 0 | immature | FM | immature |
| 03/10/2016 | 10 | 105 | 1 | immature | FM | immature |
| 03/10/2016 | 10 | 111 | 1 | milt | FM | mature |
| 03/10/2016 | 10 | 107 | 1 | immature | FM | immature |
| 03/10/2016 | 10 | 107 | 1 | milt | FM | mature |
| 03/10/2016 | 10 | 115 | 1 | milt | FM | mature |
| 03/10/2016 | 10 | 119 | 1 | milt | FM | mature |
| 03/10/2016 | 10 | 65 | 0 | immature | FM | immature |
| 03/10/2016 | 10 | 128 | 1 | immature | FM | immature |
| 03/10/2016 | 10 | 117 | 1 | milt | FM | mature |
| 03/10/2016 | 11 | 112 | 1 | milt | FM | mature |
| 03/10/2016 | 11 | 123 | 1 | milt | FM | mature |
| 03/10/2016 | 11 | 122 | 1 | mature | FM | immature |
| 03/10/2016 | 11 | 115 | 1 | immature | FM | immature |
| 03/10/2016 | 11 | 109 | 1 | milt | FM | mature |
| 03/10/2016 | 11 | 74 | 0 | immature | FM | immature |
| 03/10/2016 | 11 | 68 | 0 | immature | FM | immature |
| 03/10/2016 | 11 | 108 | 1 | milt | FM | mature |
| 03/10/2016 | 11 | 67 | 0 | immature | FM | immature |
| 03/10/2016 | 11 | 80 | 0 | milt | FM | mature |
| 03/10/2016 | 11 | 115 | 1 | immature | FM | immature |
| 03/10/2016 | 11 | 104 | 1 | milt | FM | mature |
| 03/10/2016 | 11 | 118 | 1 | immature | FM | immature |
| 03/10/2016 | 11 | 103 | 1 | immature | FM | immature |
| 03/10/2016 | 11 | 94 | 1 | milt | FM | mature |
| 03/10/2016 | 11 | 103 | 1 | immature | FM | immature |
| 03/10/2016 | 11 | 69 | 0 | immature | FM | immature |
| 03/10/2016 | 11 | 68 | 0 | immature | FM | immature |
| 03/10/2016 | 11 | 68 | 0 | immature | FM | immature |
| 03/10/2016 | 11 | 124 | 1 | immature | FM | immature |
| 03/10/2016 | 11 | 97 | 1 | milt | FM | mature |
| 03/10/2016 | 11 | 106 | 1 | milt | FM | mature |
| 03/10/2016 | 11 | 72 | 0 | immature | FM | immature |
| 03/10/2016 | 11 | 109 | 1 | milt | FM | mature |
| 03/10/2016 | 11 | 113 | 1 | immature | FM | immature |
| 03/10/2016 | 11 | 66 | 0 | immature | FM | immature |
| 03/10/2016 | 11 | 101 | 1 | milt | FM | mature |
| 03/10/2016 | 11 | 111 | 1 | immature | FM | immature |
| 03/10/2016 | 11 | 106 | 1 | mature | FM | mature |
| 03/10/2016 | 11 | 106 | 1 | immature | FM | immature |
| 03/10/2016 | 12 | 102 | 1 | immature | FM | immature |
| 03/10/2016 | 12 | 115 | 1 | immature | FM | immature |
| 03/10/2016 | 12 | 66 | 0 | immature | FM | immature |
| 03/10/2016 | 12 | 119 | 1 | immature | FM | immature |
| 03/10/2016 | 12 | 66 | 0 | immature | FM | immature |
| 03/10/2016 | 12 | 69 | 0 | immature | FM | immature |
| 03/10/2016 | 12 | 67 | 0 | immature | FM | immature |
| 03/10/2016 | 12 | 99 | 1 | mature | FM | mature |
| 03/10/2016 | 12 | 104 | 1 | milt | FM | mature |
| 03/10/2016 | 12 | 111 | 1 | milt | FM | mature |
| 03/10/2016 | 12 | 105 | 1 | immature | FM | immature |
| 03/10/2016 | 12 | 70 | 0 | immature | FM | immature |
| 03/10/2016 | 12 | 64 | 0 | immature | FM | immature |
| 03/10/2016 | 12 | 67 | 0 | immature | FM | immature |
| 03/10/2016 | 12 | 111 | 1 | milt | FM | mature |
| 03/10/2016 | 12 | 96 | 1 | milt | FM | mature |
| 03/10/2016 | 12 | 119 | 1 | milt | FM | mature |
| 03/10/2016 | 12 | 106 | 1 | milt | FM | mature |
| 03/10/2016 | 12 | 128 | 1 | milt | FM | mature |
| 03/10/2016 | 12 | 67 | 0 | immature | FM | mature |
| 03/10/2016 | 12 | 116 | 1 | milt | FM | mature |
| 03/10/2016 | 12 | 112 | 1 | milt | FM | mature |
| 03/10/2016 | 12 | 71 | 0 | immature | FM | immature |
| 03/10/2016 | 12 | 100 | 1 | milt | FM | mature |
| 03/10/2016 | 12 | 68 | 0 | immature | FM | immature |
| 03/10/2016 | 12 | 107 | 1 | immature | FM | immature |
| 03/10/2016 | 12 | 69 | 0 | immature | FM | immature |
| 03/10/2016 | 12 | 67 | 0 | immature | FM | immature |
| 03/10/2016 | 12 | 64 | 0 | immature | FM | immature |
| 03/10/2016 | 12 | 75 | 0 | immature | FM | immature |
| 03/10/2016 | 12 | 75 | 0 | milt | FM | mature |
| 03/10/2016 | 12 | 106 | 1 | milt | FM | mature |
| 03/10/2016 | 12 | 72 | 0 | immature | FM | immature |
| 03/10/2016 | 12 | 106 | 1 | immature | FM | immature |
| 03/10/2016 | 12 | 102 | 1 | immature | FM | immature |
| 03/10/2016 | 12 | 74 | 0 | immature | FM | immature |
| 03/10/2016 | 12 | 81 | 0 | milt | FM | mature |
| 03/10/2016 | 12 | 71 | 0 | immature | FM | immature |
| 03/10/2016 | 12 | 110 | 1 | immature | FM | immature |
| 03/10/2016 | 12 | 124 | 1 | milt | FM | mature |
| 03/10/2016 | 12 | 107 | 1 | milt | FM | mature |
| 03/10/2016 | 12 | 95 | 1 | milt | FM | mature |
| 03/10/2016 | 12 | 100 | 1 | milt | FM | mature |
| 03/10/2016 | 12 | 71 | 0 | immature | FM | immature |
| 03/10/2016 | 12 | 69 | 0 | immature | FM | immature |
| 03/10/2016 | 12 | 68 | 0 | immature | FM | immature |
| 03/10/2016 | 12 | 64 | 0 | immature | FM | immature |
| 03/10/2016 | 13 | 68 | 0 | immature | FM | immature |
| 03/10/2016 | 13 | 80 | 0 | immature | FM | immature |
| 03/10/2016 | 13 | 118 | 1 | milt | FM | mature |
| 03/10/2016 | 13 | 117 | 1 | immature | FM | immature |
| 03/10/2016 | 13 | 118 | 1 | milt | FM | mature |
| 03/10/2016 | 13 | 118 | 1 | immature | FM | immature |
| 03/10/2016 | 13 | 64 | 0 | immature | FM | immature |
| 03/10/2016 | 13 | 95 | 1 | milt | FM | mature |
| 03/10/2016 | 13 | 108 | 1 | immature | FM | immature |
| 03/10/2016 | 13 | 109 | 1 | immature | FM | immature |
| 03/10/2016 | 13 | 70 | 0 | immature | FM | immature |
| 03/10/2016 | 13 | 73 | 0 | immature | FM | immature |
| 03/10/2016 | 13 | 71 | 0 | immature | FM | immature |
| 03/10/2016 | 13 | 69 | 0 | immature | FM | immature |
| 03/10/2016 | 13 | 71 | 0 | immature | FM | immature |
| 03/10/2016 | 13 | 67 | 0 | immature | FM | immature |
| 03/10/2016 | 13 | 70 | 0 | immature | FM | immature |
| 03/10/2016 | 13 | 62 | 0 | immature | FM | immature |
| 03/10/2016 | 14 | 70 | 0 | immature | FM | immature |
| 03/10/2016 | 14 | 110 | 1 | immature | FM | immature |
| 03/10/2016 | 14 | 105 | 1 | milt | FM | mature |
| 03/10/2016 | 14 | 116 | 1 | immature | FM | immature |
| 03/10/2016 | 14 | 103 | 1 | milt | FM | mature |
| 03/10/2016 | 14 | 82 | 0 | immature | FM | immature |
| 03/10/2016 | 14 | 83 | 0 | immature | FM | immature |
| 03/10/2016 | 14 | 68 | 0 | immature | FM | immature |
| 03/10/2016 | 14 | 74 | 0 | immature | FM | immature |
| 03/10/2016 | 14 | 62 | 0 | immature | FM | immature |
| 03/10/2016 | 14 | 67 | 0 | immature | FM | immature |
| 03/10/2016 | 14 | 80 | 0 | immature | FM | immature |
| 03/10/2016 | 14 | 92 | 1 | milt | FM | mature |
| 03/10/2016 | 14 | 113 | 1 | milt | FM | mature |
| 03/10/2016 | 14 | 98 | 1 | milt | FM | mature |
| 03/10/2016 | 14 | 102 | 1 | mature | FM | mature |
| 03/10/2016 | 14 | 119 | 1 | immature | FM | immature |
| 03/10/2016 | 14 | 55 | 0 | immature | FM | immature |
| 03/10/2016 | 14 | 63 | 0 | immature | FM | immature |
| 03/10/2016 | 14 | 67 | 0 | immature | FM | immature |
| 03/10/2016 | 14 | 71 | 0 | immature | FM | immature |
| 03/10/2016 | 14 | 77 | 0 | immature | FM | immature |
| 03/10/2016 | 14 | 73 | 0 | immature | FM | immature |
| 03/10/2016 | 14 | 105 | 1 | immature | FM | immature |
| 03/10/2016 | 14 | 125 | 1 | milt | FM | mature |
| 03/10/2016 | 14 | 70 | 0 | immature | FM | immature |
| 03/10/2016 | 14 | 66 | 0 | immature | FM | immature |
| 03/10/2016 | 14 | 66 | 0 | immature | FM | immature |
| 03/10/2016 | 14 | 68 | 0 | immature | FM | immature |
| 03/10/2016 | 14 | 97 | 1 | milt | FM | mature |
| 03/10/2016 | 14 | 59 | 0 | immature | FM | immature |
| 03/10/2016 | 14 | 100 | 1 | mature | FM | mature |
| 03/10/2016 | 14 | 82 | 0 | mature | FM | mature |
| 03/10/2016 | 14 | 69 | 0 | immature | FM | immature |
| 03/10/2016 | 14 | 62 | 0 | immature | FM | immature |
| 03/10/2016 | 14 | 114 | 1 | milt | FM | mature |
| 03/10/2016 | 14 | 72 | 0 | mature | FM | mature |
| 03/10/2016 | 14 | 71 | 0 | milt | FM | mature |
| 03/10/2016 | 14 | 129 | 1 | milt | FM | mature |
| 03/10/2016 | 14 | 78 | 0 | immature | FM | immature |
| 03/10/2016 | 14 | 69 | 0 | mature | FM | mature |
| 03/10/2016 | 14 | 72 | 0 | immature | FM | immature |
| 03/10/2016 | 14 | 79 | 0 | immature | FM | immature |
| 03/10/2016 | 14 | 72 | 0 | immature | FM | immature |
| 03/10/2016 | 14 | 83 | 0 | immature | FM | immature |
| 03/10/2016 | 14 | 71 | 0 | immature | FM | mature |
| 03/10/2016 | 14 | 68 | 0 | immature | FM | immature |
| 03/10/2016 | 14 | 72 | 0 | immature | FM | immature |
| 03/10/2016 | 15 | 67 | 0 | immature | FM | immature |
| 03/10/2016 | 15 | 113 | 1 | mature | FM | mature |
| 03/10/2016 | 15 | 76 | 0 | immature | FM | immature |
| 03/10/2016 | 15 | 69 | 0 | immature | FM | mature |
| 03/10/2016 | 15 | 69 | 0 | mature | FM | mature |
| 03/10/2016 | 15 | 63 | 0 | immature | FM | immature |
| 03/10/2016 | 15 | 106 | 1 | milt | FM | mature |
| 03/10/2016 | 15 | 102 | 1 | immature | FM | immature |
| 03/10/2016 | 15 | 75 | 0 | immature | FM | immature |
| 03/10/2016 | 15 | 66 | 0 | immature | FM | immature |
| 03/10/2016 | 15 | 72 | 0 | immature | FM | immature |
| 03/10/2016 | 15 | 122 | 1 | milt | FM | mature |
| 03/10/2016 | 15 | 116 | 1 | milt | FM | mature |
| 03/10/2016 | 15 | 104 | 1 | milt | FM | mature |
| 03/10/2016 | 15 | 104 | 1 | milt | FM | mature |
| 03/10/2016 | 15 | 107 | 1 | milt | FM | mature |
| 03/10/2016 | 15 | 79 | 0 | immature | FM | immature |
| 03/10/2016 | 15 | 94 | 0 | immature | FM | immature |
| 03/10/2016 | 15 | 112 | 1 | immature | FM | immature |
| 03/10/2016 | 15 | 117 | 1 | immature | FM | immature |
| 03/10/2016 | 15 | 67 | 0 | immature | FM | immature |
| 03/10/2016 | 15 | 73 | 0 | immature | FM | immature |
| 03/10/2016 | 15 | 68 | 0 | immature | FM | immature |
| 03/10/2016 | 15 | 74 | 0 | immature | FM | immature |
| 03/10/2016 | 15 | 69 | 0 | immature | FM | immature |
| 03/10/2016 | 15 | 131 | 1 | milt | FM | mature |
| 03/10/2016 | 15 | 112 | 1 | milt | FM | mature |
| 03/10/2016 | 15 | 119 | 1 | milt | FM | mature |
| 03/10/2016 | 15 | 81 | 0 | immature | FM | immature |
| 03/10/2016 | 15 | 118 | 1 | immature | FM | immature |
| 03/10/2016 | 15 | 108 | 1 | milt | FM | mature |
| 03/10/2016 | 15 | 109 | 1 | milt | FM | mature |
| 03/10/2016 | 15 | 113 | 1 | immature | FM | immature |
| 03/10/2016 | 15 | 109 | 1 | immature | FM | immature |
| 03/10/2016 | 16 | 91 | 0 | immature | FM | immature |
| 03/10/2016 | 16 | 82 | 0 | immature | FM | immature |
| 03/10/2016 | 16 | 79 | 0 | milt | FM | mature |
| 04/10/2016 | 08 | 82 | 0 | immature | FM | immature |
| 04/10/2016 | 08 | 126 | 1 | milt | FM | mature |
| 04/10/2016 | 08 | 113 | 1 | immature | FM | immature |
| 04/10/2016 | 08 | 116 | 1 | immature | FM | mature |
| 04/10/2016 | 08 | 83 | 0 | immature | FM | immature |
| 04/10/2016 | 09 | 84 | 0 | immature | FM | immature |
| 04/10/2016 | 09 | 111 | 1 | milt | FM | mature |
| 04/10/2016 | 09 | 81 | 0 | milt | FM | mature |
| 04/10/2016 | 09 | 83 | 0 | mature | FM | mature |
| 04/10/2016 | 09 | 128 | 1 | immature | FM | immature |
| 04/10/2016 | 09 | 133 | 1 | immature | FM | immature |
| 04/10/2016 | 09 | 88 | 0 | immature | FM | immature |
| 04/10/2016 | 09 | 115 | 1 | milt | FM | mature |
| 04/10/2016 | 09 | 124 | 1 | milt | FM | mature |
| 04/10/2016 | 09 | 79 | 0 | milt | FM | mature |
| 04/10/2016 | 09 | 94 | 0 | immature | FM | immature |
| 04/10/2016 | 09 | 106 | 1 | milt | FM | mature |
| 04/10/2016 | 09 | 120 | 1 | milt | FM | mature |
| 04/10/2016 | 09 | 118 | 1 | immature | FM | immature |
| 04/10/2016 | 09 | 113 | 1 | milt | FM | mature |
| 04/10/2016 | 09 | 104 | 1 | milt | FM | mature |
| 04/10/2016 | 09 | 106 | 1 | milt | FM | mature |
| 04/10/2016 | 09 | 95 | 1 | milt | FM | mature |
| 04/10/2016 | 10 | 74 | 0 | mature | FM | immature |
| 04/10/2016 | 10 | 112 | 1 | milt | FM | mature |
| 04/10/2016 | 10 | 80 | 0 | immature | FM | immature |
| 04/10/2016 | 10 | 82 | 0 | immature | FM | immature |
| 04/10/2016 | 10 | 73 | 0 | immature | FM | immature |
| 04/10/2016 | 10 | 74 | 0 | immature | FM | immature |
| 04/10/2016 | 10 | 115 | 1 | mature | FM | mature |
| 04/10/2016 | 10 | 73 | 0 | mature | FM | immature |
| 04/10/2016 | 10 | 74 | 0 | immature | FM | immature |
| 04/10/2016 | 10 | 65 | 0 | immature | FM | immature |
| 04/10/2016 | 10 | 122 | 1 | milt | FM | mature |
| 04/10/2016 | 10 | 105 | 1 | milt | FM | mature |
| 04/10/2016 | 10 | 99 | 1 | milt | FM | mature |
| 04/10/2016 | 10 | 113 | 1 | milt | FM | mature |
| 04/10/2016 | 10 | 111 | 1 | milt | FM | mature |
| 04/10/2016 | 10 | 114 | 1 | milt | FM | mature |
| 04/10/2016 | 10 | 75 | 0 | mature | FM | mature |
| 04/10/2016 | 10 | 79 | 0 | immature | FM | immature |
| 04/10/2016 | 10 | 115 | 1 | immature | FM | immature |
| 04/10/2016 | 10 | 89 | 0 | immature | FM | immature |
| 04/10/2016 | 10 | 81 | 0 | immature | FM | immature |
| 04/10/2016 | 10 | 69 | 0 | immature | FM | immature |
| 04/10/2016 | 10 | 110 | 1 | immature | FM | immature |
| 04/10/2016 | 10 | 128 | 1 | immature | FM | immature |
| 04/10/2016 | 10 | 97 | 1 | milt | FM | mature |
| 04/10/2016 | 10 | 109 | 1 | immature | FM | immature |
| 04/10/2016 | 10 | 104 | 1 | immature | FM | immature |
| 04/10/2016 | 10 | 73 | 0 | immature | FM | immature |
| 04/10/2016 | 10 | 109 | 1 | immature | FM | immature |
| 04/10/2016 | 10 | 112 | 1 | milt | FM | mature |
| 04/10/2016 | 10 | 112 | 1 | milt | FM | mature |
| 04/10/2016 | 10 | 112 | 1 | milt | FM | mature |
| 04/10/2016 | 10 | 100 | 1 | milt | FM | mature |
| 04/10/2016 | 10 | 112 | 1 | milt | FM | mature |
| 04/10/2016 | 10 | 108 | 1 | mature | FM | mature |
| 04/10/2016 | 10 | 110 | 1 | milt | FM | mature |
| 04/10/2016 | 10 | 110 | 1 | immature | FM | immature |
| 04/10/2016 | 10 | 97 | 1 | milt | FM | mature |
| 04/10/2016 | 10 | 71 | 0 | immature | FM | immature |
| 04/10/2016 | 10 | 72 | 0 | immature | FM | immature |
| 04/10/2016 | 10 | 108 | 1 | milt | FM | mature |
| 04/10/2016 | 10 | 114 | 1 | milt | FM | mature |
| 04/10/2016 | 11 | 119 | 1 | milt | FM | mature |
| 04/10/2016 | 11 | 77 | 0 | immature | FM | immature |
| 04/10/2016 | 11 | 129 | 1 | immature | FM | immature |
| 04/10/2016 | 11 | 108 | 1 | milt | FM | mature |
| 04/10/2016 | 12 | 106 | 1 | immature | FM | immature |
| 04/10/2016 | 12 | 118 | 1 | immature | FM | immature |
| 04/10/2016 | 12 | 110 | 1 | immature | FM | immature |
| 04/10/2016 | 12 | 70 | 0 | immature | FM | immature |
| 04/10/2016 | 12 | 114 | 1 | immature | FM | immature |
| 04/10/2016 | 12 | 112 | 1 | milt | FM | mature |
| 04/10/2016 | 12 | 104 | 1 | immature | FM | immature |
| 04/10/2016 | 12 | 106 | 1 | milt | FM | mature |
| 04/10/2016 | 12 | 113 | 1 | milt | FM | mature |
| 04/10/2016 | 12 | 102 | 1 | milt | FM | mature |
| 04/10/2016 | 12 | 63 | 0 | immature | FM | immature |
| 04/10/2016 | 12 | 74 | 0 | mature | FM | mature |
| 04/10/2016 | 12 | 101 | 1 | milt | FM | mature |
| 04/10/2016 | 12 | 75 | 0 | mature | FM | mature |
| 04/10/2016 | 12 | 69 | 0 | immature | FM | immature |
| 04/10/2016 | 12 | 115 | 1 | immature | FM | immature |
| 04/10/2016 | 13 | 107 | 1 | milt | FM | mature |
| 04/10/2016 | 13 | 99 | 1 | immature | FM | immature |
| 04/10/2016 | 13 | 73 | 0 | immature | FM | immature |
| 04/10/2016 | 13 | 96 | 1 | milt | FM | mature |
| 04/10/2016 | 13 | 114 | 1 | milt | FM | mature |
| 04/10/2016 | 13 | 95 | 1 | milt | FM | mature |
| 04/10/2016 | 13 | 73 | 0 | immature | FM | immature |
| 04/10/2016 | 13 | 107 | 1 | milt | FM | mature |
| 04/10/2016 | 13 | 71 | 0 | milt | FM | immature |
| 04/10/2016 | 13 | 111 | 1 | immature | FM | immature |
| 04/10/2016 | 13 | 111 | 1 | milt | FM | mature |
| 04/10/2016 | 13 | 113 | 1 | immature | FM | immature |
| 04/10/2016 | 13 | 106 | 1 | milt | FM | immature |
| 04/10/2016 | 13 | 105 | 1 | immature | FM | immature |
| 04/10/2016 | 13 | 100 | 1 | milt | FM | mature |
| 04/10/2016 | 13 | 69 | 0 | immature | FM | immature |
| 04/10/2016 | 13 | 110 | 1 | milt | FM | mature |
| 04/10/2016 | 13 | 111 | 1 | immature | FM | immature |
| 04/10/2016 | 13 | 119 | 1 | milt | FM | mature |
| 04/10/2016 | 13 | 115 | 1 | immature | FM | immature |
| 04/10/2016 | 13 | 98 | 1 | milt | FM | mature |
| 04/10/2016 | 14 | 120 | 1 | immature | FM | immature |
| 04/10/2016 | 14 | 93 | 1 | milt | FM | mature |
| 04/10/2016 | 14 | 118 | 1 | immature | FM | immature |
| 04/10/2016 | 14 | 101 | 1 | milt | FM | mature |
| 04/10/2016 | 14 | 114 | 1 | mature | FM | mature |
| 04/10/2016 | 14 | 122 | 1 | milt | FM | mature |
| 04/10/2016 | 14 | 105 | 1 | milt | FM | mature |
| 04/10/2016 | 14 | 99 | 1 | milt | FM | mature |
| 04/10/2016 | 14 | 117 | 1 | milt | FM | mature |
| 04/10/2016 | 14 | 112 | 1 | milt | FM | mature |
| 04/10/2016 | 14 | 110 | 1 | immature | FM | immature |
| 04/10/2016 | 14 | 110 | 1 | mature | FM | mature |
| 04/10/2016 | 14 | 106 | 1 | milt | FM | mature |
| 04/10/2016 | 14 | 102 | 1 | milt | FM | mature |
| 04/10/2016 | 14 | 107 | 1 | immature | FM | immature |
| 04/10/2016 | 14 | 124 | 1 | immature | FM | immature |
| 04/10/2016 | 14 | 75 | 0 | immature | FM | immature |
| 04/10/2016 | 14 | 121 | 1 | milt | FM | mature |
| 04/10/2016 | 14 | 114 | 1 | immature | FM | immature |
| 04/10/2016 | 14 | 117 | 1 | immature | FM | immature |
| 04/10/2016 | 14 | 109 | 1 | milt | FM | mature |
| 04/10/2016 | 14 | 114 | 1 | immature | FM | immature |
| 04/10/2016 | 14 | 105 | 1 | immature | FM | immature |
| 04/10/2016 | 14 | 110 | 1 | immature | FM | immature |
| 04/10/2016 | 14 | 114 | 1 | immature | FM | immature |
| 04/10/2016 | 14 | 102 | 1 | milt | FM | mature |
| 04/10/2016 | 14 | 111 | 1 | milt | FM | mature |
| 04/10/2016 | 14 | 109 | 1 | immature | FM | immature |
| 04/10/2016 | 14 | 121 | 1 | milt | FM | mature |
| 04/10/2016 | 14 | 122 | 1 | immature | FM | immature |
| 04/10/2016 | 15 | 114 | 1 | immature | FM | immature |
| 04/10/2016 | 15 | 102 | 1 | milt | FM | mature |
| 04/10/2016 | 15 | 127 | 1 | milt | FM | mature |
| 04/10/2016 | 15 | 114 | 1 | immature | FM | immature |
| 04/10/2016 | 15 | 100 | 1 | immature | FM | immature |
| 04/10/2016 | 15 | 104 | 1 | milt | FM | mature |
| 04/10/2016 | 15 | 95 | 1 | immature | FM | immature |
| 04/10/2016 | 15 | 113 | 1 | immature | FM | immature |
| 04/10/2016 | 15 | 111 | 1 | immature | FM | immature |
| 04/10/2016 | 15 | 113 | 1 | immature | FM | immature |
| 04/10/2016 | 15 | 105 | 1 | immature | FM | immature |
| 04/10/2016 | 15 | 119 | 1 | immature | FM | immature |
| 04/10/2016 | 15 | 104 | 1 | milt | FM | mature |
| 04/10/2016 | 15 | 106 | 1 | immature | FM | immature |
| 04/10/2016 | 15 | 104 | 1 | immature | FM | immature |
| 04/10/2016 | 15 | 106 | 1 | milt | FM | mature |
| 04/10/2016 | 15 | 113 | 1 | milt | FM | mature |
| 05/10/2016 | 08 | 112 | 1 | milt | FM | mature |
| 05/10/2016 | 08 | 102 | 1 | milt | FM | mature |
| 05/10/2016 | 08 | 102 | 1 | milt | FM | mature |
| 05/10/2016 | 08 | 110 | 1 | milt | FM | mature |
| 05/10/2016 | 08 | 114 | 1 | milt | FM | mature |
| 05/10/2016 | 08 | 99 | 1 | milt | FM | mature |
| 05/10/2016 | 08 | 104 | 1 | immature | FM | immature |
| 05/10/2016 | 08 | 110 | 1 | milt | FM | mature |
| 05/10/2016 | 08 | 110 | 1 | immature | FM | immature |
| 05/10/2016 | 08 | 105 | 1 | milt | FM | mature |
| 05/10/2016 | 08 | 120 | 1 | immature | FM | immature |
| 05/10/2016 | 08 | 124 | 1 | immature | FM | immature |
| 05/10/2016 | 09 | 120 | 1 | immature | FM | immature |
| 05/10/2016 | 09 | 113 | 1 | milt | FM | mature |
| 05/10/2016 | 09 | 110 | 1 | milt | FM | mature |
| 05/10/2016 | 09 | 92 | 1 | milt | FM | mature |
| 05/10/2016 | 09 | 112 | 1 | milt | FM | mature |
| 05/10/2016 | 09 | 107 | 1 | immature | FM | immature |
| 05/10/2016 | 09 | 119 | 1 | immature | FM | immature |
| 05/10/2016 | 09 | 108 | 1 | immature | FM | immature |
| 05/10/2016 | 09 | 125 | 1 | immature | FM | immature |
| 05/10/2016 | 10 | 123 | 1 | milt | FM | mature |
| 05/10/2016 | 10 | 127 | 1 | immature | FM | immature |
| 05/10/2016 | 10 | 127 | 1 | immature | FM | immature |
| 05/10/2016 | 10 | 120 | 1 | milt | FM | mature |
| 05/10/2016 | 10 | 115 | 1 | milt | FM | mature |
| 05/10/2016 | 10 | 125 | 1 | immature | FM | immature |
| 05/10/2016 | 10 | 123 | 1 | milt | FM | mature |
| 05/10/2016 | 10 | 121 | 1 | milt | FM | mature |
| 05/10/2016 | 10 | 115 | 1 | immature | FM | immature |
| 05/10/2016 | 11 | 115 | 1 | milt | FM | mature |
| 05/10/2016 | 11 | 119 | 1 | immature | FM | immature |
| 05/10/2016 | 11 | 101 | 1 | milt | FM | mature |
| 05/10/2016 | 11 | 121 | 1 | milt | FM | mature |
| 05/10/2016 | 11 | 112 | 1 | milt | FM | mature |
| 05/10/2016 | 11 | 129 | 1 | milt | FM | mature |
| 05/10/2016 | 11 | 113 | 1 | milt | FM | mature |
| 05/10/2016 | 11 | 121 | 1 | milt | FM | mature |
| 05/10/2016 | 11 | 125 | 1 | immature | FM | immature |
| 05/10/2016 | 11 | 115 | 1 | immature | FM | immature |
| 05/10/2016 | 11 | 125 | 1 | milt | FM | mature |
| 05/10/2016 | 11 | 107 | 1 | milt | FM | mature |
| 05/10/2016 | 11 | 114 | 1 | immature | FM | immature |
| 05/10/2016 | 11 | 117 | 1 | milt | FM | mature |
| 05/10/2016 | 11 | 128 | 1 | immature | FM | immature |
| 05/10/2016 | 11 | 117 | 1 | milt | FM | mature |
| 05/10/2016 | 11 | 114 | 1 | milt | FM | mature |
| 05/10/2016 | 11 | 124 | 1 | immature | FM | immature |
| 05/10/2016 | 11 | 123 | 1 | immature | FM | immature |
| 05/10/2016 | 11 | 121 | 1 | milt | FM | mature |
| 05/10/2016 | 11 | 108 | 1 | milt | FM | mature |
| 05/10/2016 | 11 | 113 | 1 | immature | FM | immature |
| 05/10/2016 | 11 | 113 | 1 | milt | FM | mature |
| 05/10/2016 | 11 | 109 | 1 | milt | FM | mature |
| 05/10/2016 | 11 | 109 | 1 | milt | FM | mature |
| 05/10/2016 | 11 | 129 | 1 | immature | FM | immature |
| 05/10/2016 | 11 | 112 | 1 | immature | FM | immature |
| 05/10/2016 | 11 | 111 | 1 | immature | FM | immature |
| 05/10/2016 | 11 | 107 | 1 | immature | FM | immature |
| 05/10/2016 | 11 | 118 | 1 | immature | FM | immature |
| 05/10/2016 | 11 | 113 | 1 | immature | FM | immature |
| 05/10/2016 | 11 | 101 | 1 | milt | FM | mature |
| 05/10/2016 | 11 | 123 | 1 | immature | FM | immature |
| 05/10/2016 | 11 | 114 | 1 | immature | FM | immature |
| 05/10/2016 | 11 | 106 | 1 | milt | FM | mature |
| 05/10/2016 | 11 | 120 | 1 | milt | FM | mature |
| 05/10/2016 | 11 | 111 | 1 | immature | FM | immature |
| 05/10/2016 | 11 | 111 | 1 | immature | FM | mature |
| 05/10/2016 | 11 | 117 | 1 | milt | FM | mature |
| 05/10/2016 | 11 | 109 | 1 | immature | FM | immature |
| 05/10/2016 | 11 | 109 | 1 | milt | FM | mature |
| 05/10/2016 | 11 | 122 | 1 | immature | FM | immature |
| 05/10/2016 | 11 | 114 | 1 | milt | FM | mature |
| 05/10/2016 | 11 | 114 | 1 | immature | FM | immature |
| 05/10/2016 | 11 | 113 | 1 | immature | FM | mature |
| 05/10/2016 | 12 | 118 | 1 | mature | FM | mature |
| 05/10/2016 | 12 | 111 | 1 | immature | FM | immature |
| 05/10/2016 | 12 | 117 | 1 | immature | FM | immature |
| 05/10/2016 | 12 | 107 | 1 | immature | FM | immature |
| 05/10/2016 | 12 | 113 | 1 | immature | FM | immature |
| 05/10/2016 | 12 | 107 | 1 | immature | FM | immature |
| 09/10/2017 | 09 | 99 | 0 | immature | JT | immature |
| 09/10/2017 | 09 | 111 | 1 | milt | JT | mature |
| 09/10/2017 | 09 | 92 | 0 | immature | JT | immature |
| 09/10/2017 | 09 | 74 | 0 | immature | JT | immature |
| 09/10/2017 | 09 | 97 | 0 | immature | JT | immature |
| 09/10/2017 | 09 | 92 | 0 | immature | JT | immature |
| 09/10/2017 | 09 | 66 | 0 | immature | JT | immature |
| 09/10/2017 | 09 | 109 | 0 | immature | JT | immature |
| 09/10/2017 | 09 | 75 | 0 | immature | JT | immature |
| 09/10/2017 | 10 | 75 | 0 | immature | JT | immature |
| 09/10/2017 | 10 | 89 | 0 | immature | JT | immature |
| 09/10/2017 | 10 | 78 | 0 | immature | JT | immature |
| 09/10/2017 | 10 | 61 | 0 | immature | JT | immature |
| 09/10/2017 | 10 | 85 | 0 | immature | JT | immature |
| 09/10/2017 | 10 | 73 | 0 | immature | JT | immature |
| 09/10/2017 | 10 | 91 | 0 | milt | JT | mature |
| 09/10/2017 | 10 | 73 | 0 | immature | JT | immature |
| 09/10/2017 | 10 | 101 | 0 | immature | JT | immature |
| 09/10/2017 | 10 | 87 | 0 | immature | JT | immature |
| 09/10/2017 | 10 | 78 | 0 | immature | JT | immature |
| 09/10/2017 | 10 | 101 | 0 | immature | JT | immature |
| 09/10/2017 | 10 | 76 | 0 | immature | JT | immature |
| 09/10/2017 | 10 | 65 | 0 | immature | JT | immature |
| 09/10/2017 | 10 | 70 | 0 | immature | JT | immature |
| 09/10/2017 | 10 | 95 | 0 | immature | JT | immature |
| 09/10/2017 | 10 | 78 | 0 | immature | JT | immature |
| 09/10/2017 | 10 | 67 | 0 | immature | JT | immature |
| 09/10/2017 | 10 | 93 | 0 | immature | JT | immature |
| 09/10/2017 | 10 | 71 | 0 | mature | JT | mature |
| 09/10/2017 | 10 | 76 | 0 | immature | JT | immature |
| 09/10/2017 | 10 | 76 | 0 | immature | JT | immature |
| 09/10/2017 | 10 | 90 | 0 | immature | JT | immature |
| 09/10/2017 | 10 | 100 | 0 | immature | JT | immature |
| 09/10/2017 | 10 | 98 | 0 | immature | JT | immature |
| 09/10/2017 | 10 | 73 | 0 | immature | JT | immature |
| 09/10/2017 | 10 | 69 | 0 | immature | JT | immature |
| 09/10/2017 | 10 | 87 | 0 | immature | JT | immature |
| 09/10/2017 | 11 | 95 | 0 | milt | JT | mature |
| 09/10/2017 | 11 | 74 | 0 | milt | JT | mature |
| 09/10/2017 | 11 | 93 | 0 | immature | JT | immature |
| 09/10/2017 | 11 | 82 | 0 | milt | JT | mature |
| 09/10/2017 | 11 | 107 | 0 | immature | JT | immature |
| 09/10/2017 | 11 | 129 | 1 | immature | JT | immature |
| 09/10/2017 | 11 | 129 | 1 | immature | JT | immature |
| 09/10/2017 | 11 | 98 | 0 | immature | JT | immature |
| 09/10/2017 | 11 | 66 | 0 | immature | JT | immature |
| 09/10/2017 | 11 | 82 | 0 | milt | JT | mature |
| 09/10/2017 | 11 | 71 | 0 | immature | JT | immature |
| 09/10/2017 | 11 | 83 | 0 | immature | JT | immature |
| 09/10/2017 | 11 | 97 | 0 | immature | JT | immature |
| 09/10/2017 | 11 | 69 | 0 | immature | JT | immature |
| 09/10/2017 | 11 | 61 | 0 | immature | JT | immature |
| 09/10/2017 | 11 | 81 | 0 | milt | JT | immature |
| 09/10/2017 | 11 | 74 | 0 | immature | JT | immature |
| 09/10/2017 | 11 | 61 | 0 | immature | JT | immature |
| 09/10/2017 | 11 | 71 | 0 | immature | JT | immature |
| 09/10/2017 | 11 | 75 | 0 | immature | JT | immature |
| 09/10/2017 | 11 | 70 | 0 | immature | JT | immature |
| 09/10/2017 | 11 | 70 | 0 | immature | JT | immature |
| 09/10/2017 | 11 | 72 | 0 | milt | JT | mature |
| 09/10/2017 | 11 | 64 | 0 | immature | JT | immature |
| 09/10/2017 | 11 | 85 | 0 | immature | JT | immature |
| 09/10/2017 | 11 | 81 | 0 | immature | JT | immature |
| 09/10/2017 | 11 | 75 | 0 | immature | JT | immature |
| 09/10/2017 | 11 | 87 | 0 | immature | JT | immature |
| 09/10/2017 | 11 | 88 | 0 | immature | JT | immature |
| 09/10/2017 | 11 | 125 | 1 | immature | JT | immature |
| 09/10/2017 | 11 | 88 | 0 | immature | JT | immature |
| 09/10/2017 | 11 | 76 | 0 | immature | JT | immature |
| 09/10/2017 | 12 | 73 | 0 | immature | JT | immature |
| 09/10/2017 | 12 | 75 | 0 | immature | JT | immature |
| 09/10/2017 | 12 | 73 | 0 | immature | JT | immature |
| 09/10/2017 | 12 | 124 | 1 | immature | JT | immature |
| 09/10/2017 | 12 | 131 | 1 | immature | JT | immature |
| 09/10/2017 | 12 | 85 | 0 | immature | JT | immature |
| 09/10/2017 | 12 | 86 | 0 | immature | JT | immature |
| 09/10/2017 | 12 | 122 | 1 | immature | JT | immature |
| 09/10/2017 | 12 | 79 | 0 | immature | JT | immature |
| 09/10/2017 | 12 | 67 | 0 | immature | JT | immature |
| 09/10/2017 | 12 | 86 | 0 | immature | JT | immature |
| 09/10/2017 | 12 | 69 | 0 | immature | JT | immature |
| 09/10/2017 | 12 | 86 | 0 | immature | JT | immature |
| 09/10/2017 | 12 | 71 | 0 | immature | JT | immature |
| 09/10/2017 | 12 | 80 | 0 | immature | JT | immature |
| 09/10/2017 | 12 | 87 | 0 | immature | JT | immature |
| 09/10/2017 | 12 | 105 | 1 | immature | JT | immature |
| 09/10/2017 | 12 | 115 | 1 | milt | JT | mature |
| 09/10/2017 | 12 | 79 | 0 | immature | JT | immature |
| 09/10/2017 | 12 | 91 | 0 | immature | JT | immature |
| 09/10/2017 | 12 | 73 | 0 | immature | JT | immature |
| 09/10/2017 | 12 | 91 | 0 | immature | JT | immature |
| 09/10/2017 | 12 | 84 | 0 | immature | JT | immature |
| 09/10/2017 | 12 | 81 | 0 | immature | JT | immature |
| 09/10/2017 | 12 | 71 | 0 | immature | JT | immature |
| 09/10/2017 | 12 | 91 | 0 | immature | JT | immature |
| 09/10/2017 | 12 | 88 | 0 | immature | JT | immature |
| 09/10/2017 | 12 | 77 | 0 | immature | JT | immature |
| 09/10/2017 | 12 | 71 | 0 | immature | JT | immature |
| 09/10/2017 | 12 | 88 | 0 | immature | JT | immature |
| 09/10/2017 | 12 | 71 | 0 | immature | JT | immature |
| 09/10/2017 | 12 | 75 | 0 | immature | JT | mature |
| 09/10/2017 | 12 | 73 | 0 | immature | JT | immature |
| 09/10/2017 | 12 | 81 | 0 | milt | JT | mature |
| 09/10/2017 | 13 | 100 | 0 | immature | JT | immature |
| 09/10/2017 | 13 | 62 | 0 | immature | JT | immature |
| 09/10/2017 | 13 | 100 | 0 | immature | JT | immature |
| 09/10/2017 | 13 | 86 | 0 | immature | JT | immature |
| 09/10/2017 | 13 | 89 | 0 | immature | JT | immature |
| 09/10/2017 | 13 | 89 | 0 | immature | JT | immature |
| 09/10/2017 | 13 | 87 | 0 | immature | JT | immature |
| 09/10/2017 | 13 | 67 | 0 | immature | JT | immature |
| 09/10/2017 | 13 | 72 | 0 | immature | JT | immature |
| 09/10/2017 | 13 | 76 | 0 | immature | JT | immature |
| 09/10/2017 | 13 | 76 | 0 | immature | JT | immature |
| 09/10/2017 | 13 | 66 | 0 | immature | JT | immature |
| 09/10/2017 | 13 | 122 | 1 | milt | JT | mature |
| 09/10/2017 | 13 | 119 | 1 | immature | JT | immature |
| 09/10/2017 | 13 | 108 | 1 | milt | JT | mature |
| 09/10/2017 | 13 | 112 | 1 | immature | JT | immature |
| 09/10/2017 | 14 | 122 | 1 | immature | JT | immature |
| 09/10/2017 | 14 | 90 | 0 | immature | JT | immature |
| 09/10/2017 | 14 | 123 | 1 | milt | JT | mature |
| 09/10/2017 | 14 | 71 | 0 | immature | JT | immature |
| 09/10/2017 | 14 | 74 | 0 | immature | JT | immature |
| 09/10/2017 | 14 | 99 | 0 | immature | JT | immature |
| 09/10/2017 | 14 | 69 | 0 | immature | JT | immature |
| 09/10/2017 | 14 | 84 | 0 | immature | JT | immature |
| 09/10/2017 | 14 | 120 | 1 | immature | JT | immature |
| 09/10/2017 | 14 | 83 | 0 | immature | JT | immature |
| 09/10/2017 | 14 | 96 | 0 | immature | JT | immature |
| 09/10/2017 | 14 | 56 | 0 | immature | JT | immature |
| 09/10/2017 | 14 | 73 | 0 | milt | JT | immature |
| 09/10/2017 | 14 | 80 | 0 | immature | JT | immature |
| 09/10/2017 | 15 | 104 | 0 | immature | JT | immature |
| 09/10/2017 | 15 | 122 | 1 | milt | JT | mature |
| 09/10/2017 | 15 | 77 | 0 | immature | JT | immature |
| 09/10/2017 | 15 | 118 | 1 | immature | JT | immature |
| 09/10/2017 | 15 | 115 | 1 | immature | JT | immature |
| 09/10/2017 | 15 | 96 | 0 | immature | JT | immature |
| 09/10/2017 | 15 | 107 | 1 | milt | JT | mature |
| 09/10/2017 | 15 | 82 | 0 | immature | JT | immature |
| 09/10/2017 | 15 | 97 | 0 | immature | JT | immature |
| 09/10/2017 | 15 | 91 | 0 | immature | JT | immature |
| 09/10/2017 | 15 | 85 | 0 | immature | JT | immature |
| 09/10/2017 | 16 | 89 | 0 | milt | JT | mature |
| 09/10/2017 | 16 | 93 | 0 | immature | JT | immature |
| 09/10/2017 | 16 | 74 | 0 | immature | JT | immature |
| 09/10/2017 | 16 | 99 | 0 | immature | JT | immature |
| 09/10/2017 | 16 | 88 | 0 | immature | JT | immature |
| 09/10/2017 | 16 | 87 | 0 | milt | JT | mature |
| 09/10/2017 | 16 | 101 | 0 | immature | JT | immature |
| 09/10/2017 | 16 | 95 | 0 | immature | JT | immature |
| 09/10/2017 | 16 | 93 | 0 | immature | JT | immature |
| 09/10/2017 | 16 | 93 | 0 | immature | JT | immature |
| 09/10/2017 | 16 | 75 | 0 | immature | JT | mature |
| 10/10/2017 | 09 | 96 | 0 | immature | JT | immature |
| 10/10/2017 | 09 | 128 | 1 | milt | JT | mature |
| 10/10/2017 | 09 | 96 | 0 | immature | JT | immature |
| 10/10/2017 | 09 | 100 | 1 | immature | JT | immature |
| 10/10/2017 | 10 | 120 | 1 | milt | JT | immature |
| 10/10/2017 | 10 | 85 | 0 | immature | JT | immature |
| 10/10/2017 | 10 | 79 | 0 | immature | JT | mature |
| 10/10/2017 | 10 | 81 | 0 | immature | JT | immature |
| 10/10/2017 | 10 | 106 | 0 | immature | JT | immature |
| 10/10/2017 | 10 | 86 | 0 | milt | JT | mature |
| 10/10/2017 | 10 | 91 | 0 | immature | JT | immature |
| 10/10/2017 | 10 | 100 | 0 | immature | JT | immature |
| 10/10/2017 | 10 | 95 | 0 | immature | JT | immature |
| 10/10/2017 | 10 | 90 | 0 | immature | JT | immature |
| 10/10/2017 | 10 | 85 | 0 | immature | JT | immature |
| 10/10/2017 | 10 | 113 | 1 | immature | JT | immature |
| 10/10/2017 | 10 | 70 | 0 | immature | JT | immature |
| 10/10/2017 | 10 | 84 | 0 | milt | JT | mature |
| 10/10/2017 | 11 | 90 | 0 | immature | JT | immature |
| 10/10/2017 | 11 | 104 | 0 | immature | JT | immature |
| 10/10/2017 | 11 | 90 | 0 | immature | JT | immature |
| 10/10/2017 | 11 | 104 | 0 | immature | JT | immature |
| 10/10/2017 | 11 | 86 | 0 | immature | JT | immature |
| 10/10/2017 | 11 | 82 | 0 | immature | JT | immature |
| 10/10/2017 | 11 | 87 | 0 | immature | JT | immature |
| 10/10/2017 | 11 | 126 | 1 | immature | JT | immature |
| 10/10/2017 | 11 | 95 | 0 | immature | JT | immature |
| 10/10/2017 | 11 | 114 | 1 | milt | JT | mature |
| 10/10/2017 | 11 | 92 | 0 | immature | JT | immature |
| 10/10/2017 | 11 | 74 | 0 | milt | JT | mature |
| 10/10/2017 | 11 | 89 | 0 | immature | JT | immature |
| 10/10/2017 | 11 | 89 | 0 | immature | JT | immature |
| 10/10/2017 | 12 | 86 | 0 | immature | JT | immature |
| 10/10/2017 | 12 | 92 | 0 | immature | JT | immature |
| 10/10/2017 | 12 | 98 | 0 | milt | JT | mature |
| 10/10/2017 | 12 | 88 | 0 | immature | JT | immature |
| 10/10/2017 | 12 | 95 | 0 | immature | JT | immature |
| 10/10/2017 | 12 | 74 | 0 | immature | JT | immature |
| 10/10/2017 | 12 | 93 | 0 | immature | JT | immature |
| 10/10/2017 | 12 | 81 | 0 | milt | JT | mature |
| 10/10/2017 | 12 | 92 | 0 | immature | JT | immature |
| 10/10/2017 | 12 | 82 | 0 | immature | JT | immature |
| 10/10/2017 | 12 | 79 | 0 | immature | JT | immature |
| 10/10/2017 | 12 | 88 | 0 | immature | JT | immature |
| 10/10/2017 | 12 | 82 | 0 | immature | JT | immature |
| 10/10/2017 | 12 | 81 | 0 | immature | JT | immature |
| 10/10/2017 | 12 | 81 | 0 | immature | JT | immature |
| 10/10/2017 | 12 | 75 | 0 | immature | JT | immature |
| 10/10/2017 | 14 | 126 | 1 | milt | JT | immature |
| 11/10/2017 | 11 | 116 | 0 | immature | JT | immature |

## References

Baglinière, J.-L., and Maisse, G. 1985. Precocious maturation and smoltification in wild Atlantic salmon in the Armorican massif, France. Aquaculture 45(1): 249–263. doi:10.1016/0044-8486(85)90274-1.

Berejikian, B.A., Van Doornik, D.M., Endicott, R.C., Hoffnagle, T.L., Tezak, E.P., Moore, M.E., and Atkins, J. 2010. Mating success of alternative male phenotypes and evidence for frequency-dependent selection in Chinook salmon, Oncorhynchus tshawytscha. Can. J. Fish. Aquat. Sci. 67(12): 1933–1941. doi:10.1139/F10-112.

Buoro, M., Prévost, E., and Gimenez, O. 2010. Investigating Evolutionary Trade-Offs in Wild Populations of Atlantic Salmon (Salmo salar): Incorporating Detection Probabilities and Individual Heterogeneity. Evolution 64(9): 2629–2642. doi:10.1111/j.1558-5646.2010.01029.x.

Caswell, H., Naiman, R.J., and Morin, R. 1984. Evaluating the consequence of reproduction in complex salmonid life cycles. Aquaculture 43: 123–134.

Chaput, G. 2012. Overview of the status of Atlantic salmon (Salmo salar) in the North Atlantic and trends in marine mortality. ICES J. Mar. Sci. 69(9): 1538–1548. doi:10.1093/icesjms/fss013.

Clutton-Brock, T., and Sheldon, B.C. 2010. Individuals and populations: the role of long-term, individual-based studies of animals in ecology and evolutionary biology. Trends Ecol. Evol. 25(10): 562–573.

Fleming, I.A. 1996. Reproductive strategies of Atlantic salmon: ecology and evolution. Rev. Fish Biol. Fish. 6(4): 379–416. doi:10.1007/BF00164323.

Genovart, M., Pradel, R., and Oro, D. 2012. Exploiting uncertain ecological fieldwork data with multi-event capture–recapture modelling: an example with bird sex assignment. J. Anim. Ecol.: no-no. doi:10.1111/j.1365-2656.2012.01991.x.

Hutchings, J.A., and Myers, R.A. 1994. The evolution of alternative mating strategies in variable environments. Evol. Ecol. 8(3): 256–268.

ICES. 2013. Report of the Working Group on North Atlantic Salmon (WGNAS). Copenhagen, Denmark.

Irvine, J.R., and Fukuwaka, M. 2011. Pacific salmon abundance trends and climate change. ICES J. Mar. Sci. 68(6): 1122–1130. doi:10.1093/icesjms/fsq199.

Marchand, F., Azam, D., Delanoë, R., Destouches, J.-P., Tremblay, J., Baglinière, J.-L., Nevoux, M., Prévost, E., and Azam, D. 2018. Abundances and biological traits of of fish sampled by electrofishing (except specific abundance indices) on the River Oir since 1988 (France). doi:10.15468/yaikgn.

Martin, R.W., Myers, J., Sower, S.A., Phillips, D.J., and Mcauley, C. 1983. Ultrasonic Imaging, a Potential Tool for Sex Determination of Live Fish. North Am. J. Fish. Manag. 3(3): 258–264. doi:10.1577/1548-8659(1983)3<258:UIAPTF>2.0.CO;2.

Mattson, N.S. 1991. A new method to determine sex and gonad size in live fishes by using ultrasonography. J. Fish Biol. 39(5): 673–677. doi:10.1111/j.1095-8649.1991.tb04397.x.

Moran, P., Pendás, A.M., Beall, E., and García-Vázquez, E. 1996. Genetic assessment of the reproductive success of Atlantic salmon precocious parr by means of VNTR loci. Heredity 77(6): 655–660. doi:10.1038/hdy.1996.193.

Myers, R.A. 1984. Demographic Consequences of Precocious Maturation of Atlantic Salmon (Salmo salar). Can. J. Fish. Aquat. Sci. 41(9): 1349–1353. doi:10.1139/f84-165.

Myers, R.A., Hutchings, J.A., and Gibson, R.J. 1986. Variation in Male Parr Maturation Within and Among Populations of Atlantic Salmon, Salmo salar. Can. J. Fish. Aquat. Sci. 43(6): 1242–1248. doi:10.1139/f86-154.

Novelo, N.D., and Tiersch, T.R. 2012. A Review of the Use of Ultrasonography in Fish Reproduction. North Am. J. Aquac. 74(2): 169–181. doi:10.1080/15222055.2012.672370.

Parmesan, C. 2006. Ecological and evolutionary responses to recent climate change. Annu. Rev. Ecol. Evol. Syst. 37(1): 637–669.

R Development Core Team. 2018. R: A language and environment for statistical computing. R Foundation for Statistical Computing, Vienna, Austria. Available from http://www.R-project.org.

